# Hostile attribution bias shapes neural synchrony in the left ventromedial prefrontal cortex during ambiguous social narratives

**DOI:** 10.1101/2023.07.11.548407

**Authors:** Yizhou Lyu, Zishan Su, Dawn Neumann, Kimberly L. Meidenbauer, Yuan Chang Leong

**Author notes:** These authors contributed equally to this work (Yizhou Lyu, Zishan Su).

## Abstract

Hostile attribution bias refers to the tendency to interpret social situations as intentionally hostile. While previous research has focused on its developmental origins and behavioral consequences, the underlying neural mechanisms remain underexplored. Here, we employed functional near infrared spectroscopy (fNIRS) to investigate the neural correlates of hostile attribution bias. While undergoing fNIRS, participants listened to and provided attribution ratings for 21 hypothetical scenarios where a character’s actions resulted in a negative outcome for the listener. Ratings of hostile intentions were averaged to obtain a measure of hostile attribution bias. Using intersubject-representational similarity analysis, we found that participants with similar levels of hostile attribution bias exhibited higher levels of neural synchrony during narrative listening, suggesting shared interpretations of the scenarios. This effect was localized to the left ventromedial prefrontal cortex (VMPFC), and was particularly prominent in scenarios where the character’s intentions were highly ambiguous. We then grouped participants into high and low bias groups based on a median split of their hostile attribution bias scores. A similarity-based classifier trained on the neural data classified participants as having high or low bias with 76% accuracy, indicating that the neural time courses during narrative listening was systematically different between the two groups. Furthermore, hostile attribution bias correlated negatively with attributional complexity, a measure of one’s tendency to consider multifaceted causes when explaining behavior. Our study sheds light on the neural mechanisms underlying hostile attribution bias and highlights the potential of using fNIRS to develop non-intrusive and cost-effective neural markers of this socio-cognitive bias.

**Significance Statement:** Inferring the intentions from behavior is crucial for adaptive social functioning. A predisposition towards interpreting intentions as hostile is a significant predictor of interpersonal conflict and aggressive tendencies. Using fNIRS, we found that individual differences in hostile attribution bias shaped neural synchrony in the VMPFC while processing real-world social situations. Additionally, we were able to distinguish between participants with high and low hostile attribution bias from neural activity time courses. These results reveal how subjective interpretations of social situations are influenced by hostile attribution bias and reflected in the temporal dynamics of the VMPFC. Our findings lay the groundwork for future studies aimed at understanding the neurobiological basis of socio-cognitive biases, as well as interventions aimed at mitigating these biases.

## Introduction

Picture two strangers walking down a crowded sidewalk, brushing shoulders as they pass. While one of them might view this as an innocuous and unavoidable outcome of the crowded environment, the other might perceive it as a deliberate hostile act. Hostile attribution bias refers to a predisposition to perceive others’ actions as hostile, and is thought to reflect a skewed system of appraisals and expectancies that biases social judgments (Epps and Kendall, 1995; Dodge, 2006; Klein Tuente et al., 2019). The tendency to infer hostile intentions predisposes an individual to respond aggressively, resulting in a self-fulfilling prophecy where perceived hostility begets actual hostility. Indeed, hostile attribution bias has been tightly linked to increased physical and relational aggression, impaired social relationships, and poor mental health (Pettit et al., 2010; Dodge et al., 2015; Smith et al., 2016). Studying the neural basis of hostile attribution bias can inform targeted interventions that reduce aggressive behavior and foster healthier relationships.

While the implications of hostile attribution bias on maladaptive behavior are extensively documented, its neurobiological underpinnings remain relatively underexplored. Prior studies have used structural MRI and lesion-mapping methods to identify brain regions associated with individual differences in hostile attribution bias and aggression (Grafman et al., 1996; Yang and Raine, 2009; Cristofori et al., 2016; Quan et al., 2019). A consistent finding emerging from these studies is that structural differences (e.g. morphological variation or lesions) in the prefrontal cortex is associated with the tendency to make hostile attributions and behave aggressively. For example, Quan and colleagues (2019) found that larger gray matter volume in the left orbitofrontal cortex (OFC) was associated with higher trait-level hostile attribution bias. Furthermore, gray matter volume in the left OFC mediated the effects of hostile attribution bias on participants’ willingness to endorse violence. In line with these findings, non-invasive stimulation of the prefrontal cortex modulates aggressive tendencies (Hortensius et al., 2012; Dambacher et al., 2015; Choy et al., 2018).

The prior work suggests a pivotal role of the prefrontal cortex in the manifestation and regulation of hostile attribution bias. These studies have examined the structural correlates of hostile attribution bias and its downstream behavioral effects, but it remains unclear if and how prefrontal regions are engaged during the ongoing processing of social information that ultimately gives rise to hostile attributions. Furthermore, there is a lack of spatial specificity in the prefrontal regions involved, with some studies highlighting ventromedial regions (Grafman et al., 1996; Quan et al., 2019), and others highlighting dorsolateral regions (Cristofori et al., 2016; Choy et al., 2018). Additionally, existing functional neuroimaging studies have focused on aggression and violence (Yang and Raine, 2009; Fanning et al., 2017), and not the attributional biases that result in the hostile interpretation of social cues.

The goal of this study is to investigate the dynamic engagement of prefrontal regions during real-time processing of social situations and how they contribute to hostile attribution bias. We used functional near-infrared spectroscopy (fNIRS) to measure activity in the prefrontal cortex as participants listened to audio narratives that were created to measure hostile attribution bias (Epps and Kendall, 1995). These scenarios described nuanced social situations where a character acted in a manner that led to a negative outcome for the listener. Hostile attribution bias was measured by evaluating participants’ hostility ratings of the characters’ actions. Due to the mixed evidence implicating different prefrontal regions, we measured activity across the prefrontal cortex. We then used intersubject-representational similarity analysis (Finn et al., 2020) to test the hypothesis that individuals with similar levels of hostile attribution bias would have similar neural dynamics while listening to the narratives, reflecting similar interpretations of the scenarios. Thus, our paradigm allowed us to relate activity of specific prefrontal regions during ongoing processing of social situations to individual differences in the tendency to attribute hostile intentions, paving the way for a deeper understanding of the neurobiological mechanisms underpinning hostile attribution bias.

## Materials and Methods

### Participants

Sixty-four individuals participated in the study. Experimental procedures were approved by the University of Chicago Institutional Review Board, and all participants provided informed consent prior to the start of the study. Participants received either 1 course credit or $15 for the hour-long experiment. All participants self-reported having native proficiency in English, no hearing or speech comprehension disorders, no serious head injury, no neurological/psychiatric disorders, and were not taking psychiatric medication. Individuals were recruited from the University of Chicago community through the research participation system managed by the Department of Psychology (SONA systems). The study advertisement and consent form indicated that the study was an investigation of the neural basis of how people understand narratives. Four participants had incomplete data due to technical difficulties with the recording device and were excluded from analyses. Additionally, data from two participants were excluded because of unusable data (see fNIRS data acquisition and preprocessing), yielding an effective sample size of fifty-eight participants (Table 1). All participants who completed the study reported that they were able to comprehend the narratives.

**Table 1.**
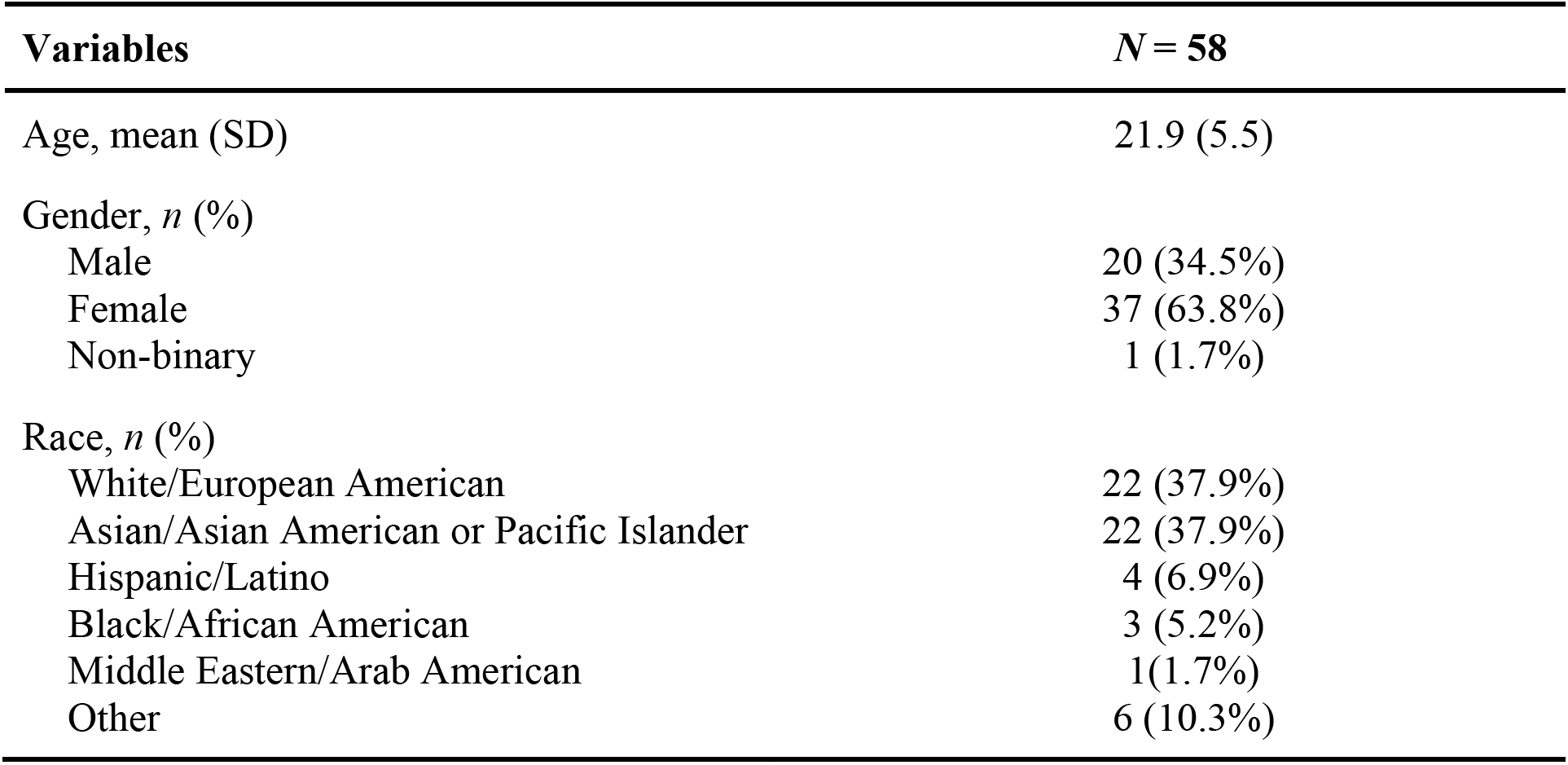
Participant demographics

| **Variables** | ***N* = 58** |
| --- | --- |
| Age, mean (SD) | 21.9 (5.5) |
| Gender, *n* (%) |  |
| Male | 20 (34.5%) |
| Female | 37 (63.8%) |
| Non-binary | 1 (1.7%) |
| Race, *n* (%) |  |
| White/European American | 22 (37.9%) |
| Asian/Asian American or Pacific Islander | 22 (37.9%) |
| Hispanic/Latino | 4 (6.9%) |
| Black/African American | 3 (5.2%) |
| Middle Eastern/Arab American | 1 (1.7%) |
| Other | 6 (10.3%) |

### Experimental Procedures

Participants were scanned using fNIRS while they listened to 21 narrated scenarios (total duration: 13 min 30 s; average duration: 38.57s; Fig. 1A). These scenarios were taken from a scenario-based questionnaire used to measure hostile attribution bias in adult samples (Epps and Kendall, 1995). In each scenario, the character in the story acted in a way that resulted in a hypothetical negative outcome for the listener (e.g., a former employer forgetting to submit a letter of recommendation). Prior work conducted by the author of these scenarios shows that, on average, they elicit hostile, benign and ambiguous attributions of the character’s intentions (hostile, benign and ambiguous scenarios; seven in each category). We used this questionnaire over others due to the relatively large number of scenarios and because the scenarios tended to be longer and described nuanced social situations. We recorded a research assistant narrating each scenario in a neutral tone. The use of audio rather than written stimuli allowed for more precise temporal control over how participants processed the scenarios.

**Figure 1.**
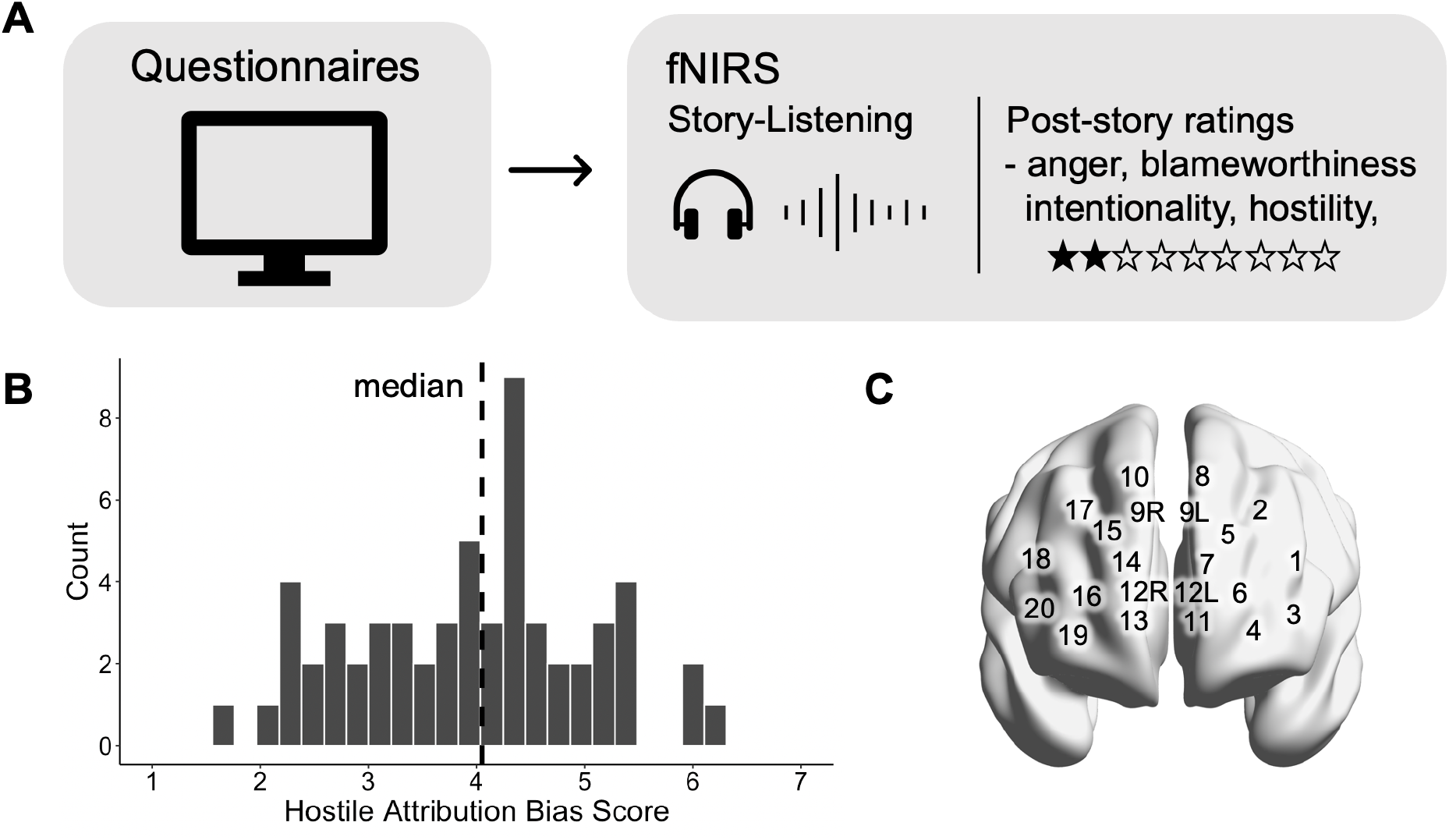
Experimental design. **(A)** Prior to the experiment, participants completed a series of behavioral questionnaires that measured individual differences in aggression, social connectedness and attributional complexity. They then listened to 21 narrated social scenarios while undergoing fNIRS. At the end of each story, participants provided hostility, intentionality, blameworthiness and anger ratings. **(B)** We computed each participant’s hostile attribution bias score as the average hostility ratings across all scenarios. Dotted line indicates sample median. **(C)** Spatial location of the 20 fNIRS channels projected onto the cortical surface.

While listening, participants were asked to imagine and react to the scenarios as if they were happening to them. Following each story, participants rated how well they understood the recording (comprehension ratings), how angry they would be if the incident happened to them (anger ratings), if they thought the character intended their actions (intentionality ratings), if they thought the person’s actions were hostile (hostility ratings), and the extent to which they blamed the character for their actions (blameworthiness ratings). All ratings were collected from a 0 to 9 point scale, and rescaled to a 1 to 9 point scale to facilitate comparison with prior studies that have used these scenarios (e.g., Epps and Kendall, 1995; Neumann et al., 2015, 2021).

### fNIRS data acquisition and preprocessing

fNIRS data were collected using a NIRSport2 fNIRS unit (NIRx Medical Technologies) with a layout of 20 channels composed of 8 light sources and 7 detectors. Spatial positions were standardized across participants using the unambiguously illustrated (UI) 10/10 external positioning system (Jurcak et al., 2007). Data were collected at a sampling rate of 10.1725Hz at wavelengths of 760 nm and 850 nm. Due to a limited number of available optodes, it was not possible to obtain whole brain fNIRS coverage. Optodes were placed to optimize coverage of the prefrontal cortex (PFC) due to extensive prior work implicating the region in aggression (Harmon-Jones and Sigelman, 2001; Yang and Raine, 2009; Cristofori et al., 2016; Choy et al., 2018), social attributions (Forbes and Grafman, 2010; Spunt and Lieberman, 2012; Wagner et al., 2019; Elder et al., 2023), and the subjective interpretation of social narratives (Yeshurun et al., 2017; Finn et al., 2018; Nguyen et al., 2019; Leong et al., 2020; Dieffenbach et al., 2021), suggesting that the region may play a critical role in the manifestation and regulation of hostile attribution bias.

Data preprocessing was performed using custom MATLAB scripts that utilized the Homer2 analysis package (Huppert et al., 2009). Channels were identified as unusable and removed if detector saturation occurred for longer than two seconds, or if the signal’s power spectrum exceeded a quartile coefficient of dispersion of 0.29 over the course of the scan (Dieffenbach et al., 2021). Data from two participants were discarded for having more than 50% unusable channels. For each channel, the raw light intensity was converted to optical density, and a bandpass filter of 0.005 to 0.5Hz was applied to remove the influence of cardiac activity and low-frequency drift. Motion artifacts were identified and corrected using targeted principal components analysis as implemented in Homer2 (Yücel et al., 2014). Specifically, motion artifacts were identified as optical density values that exceeded five standard deviations or an absolute change in signal amplitude of two within a one-second interval. A principal components analysis was then performed to remove 80% of the variance in the one-second around the identified artifact.

Optical density values were then converted to changes in oxygenated (HbO), deoxygenated (HbR) and total (HbT) hemoglobin concentrations using the Modified Beer Lambert Law. All analyses utilized the HbO signal, which has been shown to have stronger signal amplitude, higher correlation to fMRI BOLD signals, and better signal-to-noise ratio than HbR and HbT (Strangman et al., 2002; Tong and Frederick, 2010; Duan et al., 2012). Finally, the HbO time course of each participant was shifted by 4.5 seconds to account for the lag in the hemodynamic response (Huppert et al., 2006) and z-scored across time.

### Intersubject-representational similarity analyses

We employed intersubject-representational similarity analysis (IS-RSA; Nguyen et al., 2019; Chen et al., 2020; Finn et al., 2020; Fig. 2A) to examine the relationship between hostile attribution bias and neural synchrony during narrative listening. IS-RSA builds on classic RSA (Kriegeskorte et al., 2008), where the similarity between different experimental conditions is compared to the similarity in neural responses to the experimental conditions. A correlation between the two would indicate a correspondence in how internal task representations are expected to differ based on the experimental conditions and the neural representation in a given brain region. Here, instead of comparing responses between experimental conditions within a participant, we compared the responses to the same experimental stimuli across participants to test whether participants with more similar levels of hostile attribution bias also exhibit greater similarity in their neural responses.

**Figure 2.**
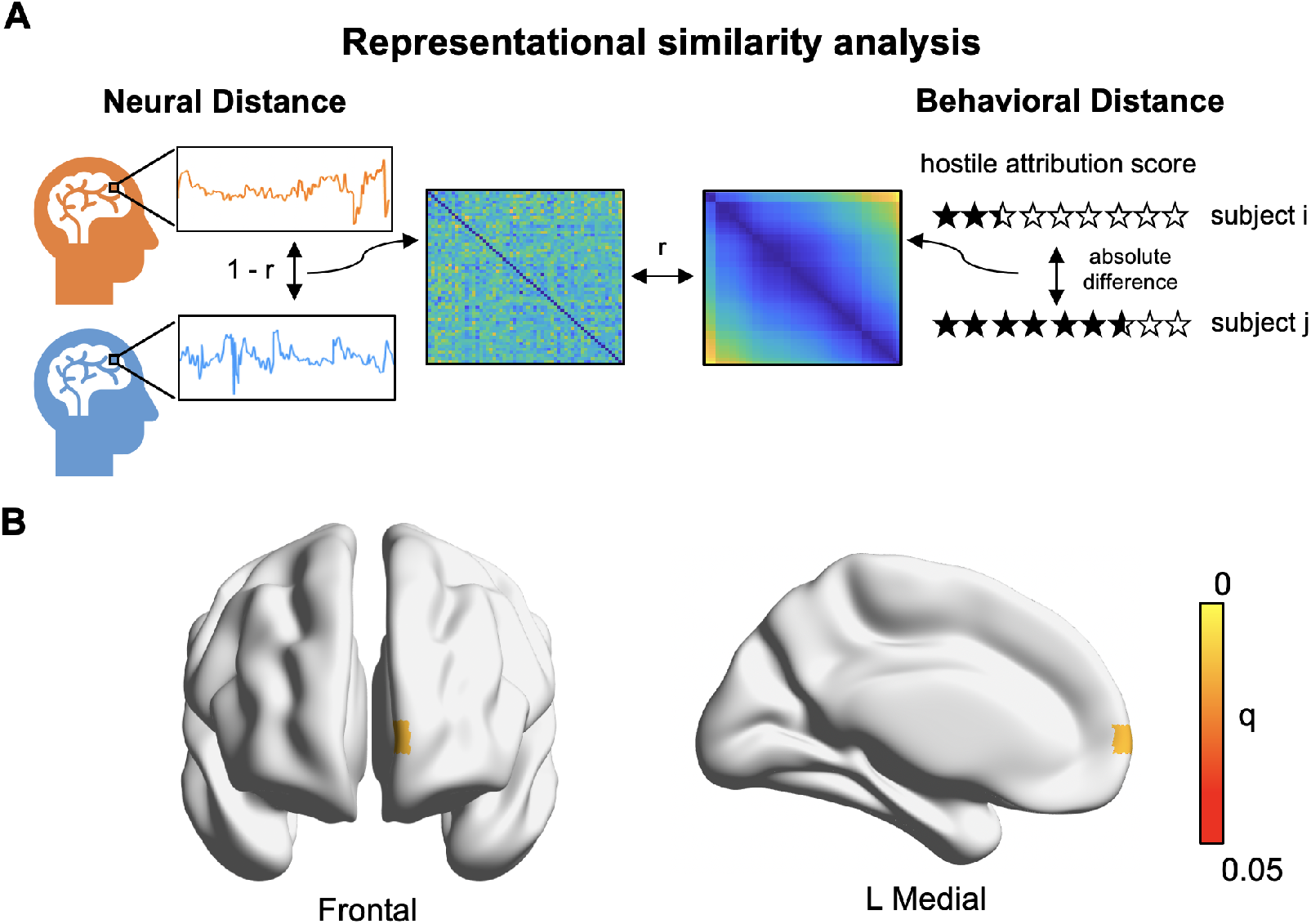
Hostile attribution bias shapes neural synchrony in the VMPFC. **(A)** Schematic of intersubject-representational similarity analysis. For each channel, we constructed a neural distance matrix by calculating the 1 - Pearson correlation between the time courses of every pair of participants. We then correlated each neural distance matrix with a behavioral distance matrix constructed from the absolute difference in hostile attribution bias scores for every pair of participants. **(B)** Left VMPFC (Channel 11) activity time course was more similar in individuals with similar levels of hostile attribution bias. *p-*values were computed nonparametrically using a Mantel test, and thresholded at an FDR of *q* < 0.05.

For each participant, we first computed their hostile attribution bias score as the average hostility rating across the 21 stories. We then constructed a *behavioral distance matrix* by calculating the absolute difference in hostile attribution scores between every pair of participants. As such, the behavioral distance matrix reflects the similarity structure in hostile attribution across participants, with smaller distances denoting pairs of participants with similar levels of hostile attribution bias. For each fNIRS channel of each participant, we extracted and concatenated the activity time course while the participant listened to the 21 narratives, and z-scored the time courses across time. For each channel, we constructed a *neural distance matrix* by computing the correlation distance (i.e. 1 - Pearson’s *r*) of the concatenated time courses between every pair of participants. The neural distance matrix reflects the similarity structure in the temporal dynamics of neural activity between pairs of participants at that corresponding channel, with smaller distances denoting pairs of participants whose activity time course was more similar. As the distance matrices were symmetrical along the diagonal, only the lower triangle of each matrix (excluding the diagonal) was retained.

We then computed the Pearson’s correlation between the behavioral distance matrix and the neural distance matrix for each channel. A positive correlation indicates a channel where activity time courses were more synchronous between participants who were similar in hostile attribution bias. Statistical significance was assessed using a Mantel test (Mantel, 1967; Kriegeskorte *et al*., 2008). Specifically, we recomputed the IS-RSA with behavioral distance matrices constructed from data where the identities of the participants were shuffled such that the distance matrix no longer reflects the similarity structure observed in the data. This procedure was repeated 10,000 times to generate a null distribution, with the p-value calculated as the proportion of the null distribution with a r-value that was more positive than the empirical r-value. We then corrected for multiple comparisons across 20 channels by controlling for the false discovery rate (FDR; Benjamini and Hochberg, 1995) of *q* < 0.05 using the *p.adjust* function implemented in base R.

To test if our results would be robust to a different metric of neural distance, we reran all analyses using neural distance matrices constructed from the Euclidean distance between the activity time courses of participant pairs. To examine the relationship between hostile attribution bias and neural synchrony for hostile, ambiguous and benign scenarios, we reran the IS-RSA for each scenario type. To examine the relationship between neural synchrony and intersubject similarity in intentionality attributions, anger and assessments of blameworthiness, we reran the IS-RSA with behavioral distance matrices constructed from intentionality, anger and blameworthiness ratings. For each behavioral measure, we first computed the average rating across the 21 scenarios for each participant. The behavioral distance matrix was then computed from the absolute difference in average ratings of the corresponding measure between each pair of participants, and correlated with the neural distance matrix.

Additionally, we tested if an *Anna Karenina* (AnnaK) model (Finn et al., 2020; Baek et al., 2023) of intersubject representational similarity would account for the neural data. The *Anna Karenina* model is named after the opening line of the Tolstoy novel: “All happy families are alike; each unhappy family is unhappy in its own way”, and captures the hypothesis that some participants might be alike, while others are idiosyncratic. We constructed an *AnnaK* behavioral distance matrix where each cell is computed as the average hostile attribution bias score of each pair of participants. In the AnnaK behavioral matrix, pairs of participants with low hostile attribution bias would have a low distance value, while pairs of participants with high hostile attribution bias would have a high distance value. We then computed the correlation between the AnnaK behavioral matrix and the neural distance matrix of each channel. A positive correlation would indicate that neural responses were more synchronized among participants with low hostile attribution bias, but were more idiosyncratic among participants with high hostile attribution bias. In contrast, a negative correlation would suggest the opposite, i.e., neural responses were more synchronized among participants with high hostile attribution bias, but were more idiosyncratic among participants with low hostile attribution bias. Statistical significance was again assessed using Mantel tests and corrected at FDR *q* < 0.05.

### Analyses on mean fNIRS activity

For each participant and channel, we computed the mean neural activity across the duration for each scenario. We then averaged the mean neural activity across scenarios separately for hostile, ambiguous and benign narratives. Two-sided paired-sample t-tests were then used to test if mean activity at each channel was different between (1) hostile and ambiguous narratives, (2) hostile and benign narratives, and (3) ambiguous and benign narratives. Next, we examined if individual differences in hostile attribution bias would moderate differences in mean activity between narrative types. For each of the three contrasts, we correlated each participant’s hostile bias score with the mean difference in activity between the two narrative types. In all analyses described in this section, we applied an FDR-correction of *q* < 0.05 separately for each of the three contrasts.

### Synchrony-based classification analysis

We adapted a synchrony-based classification approach used in prior fNIRS and fMRI studies (Yeshurun et al., 2017; Dieffenbach et al., 2021) to assess the extent to which we were able to identify participants with high and low hostile attribution bias. Participants were first grouped into high and low hostile attribution groups using a median split of their hostile attribution bias scores (n = 29 in each group). For each iteration of a leave-one-out cross validation procedure, a participant was held out as the test data. For the remaining participants, we averaged the neural time courses of each channel separately for each group to obtain a “template” time course that captured the temporal dynamics of neural activity for the respective group. The activity time course for the held-out participant was then correlated with each of the two template time courses. The participant was classified as high hostile attribution bias if the correlation was higher with the high hostile attribution bias template, and classified as low hostile attribution bias if the correlation was higher with the low hostile attribution bias template. Classification accuracy was computed separately at each channel. Statistical significance was assessed using a non-parametric permutation test by comparing the true classification accuracy to a null distribution generated by re-running the analysis 10,000 times with shuffled group labels. Results were then thresholded at an FDR *q* of < 0.05.

### Spatial localization and visualization of fNIRS channels

To spatially localize and visualize the fNIRS results, approximate MNI coordinates of each fNIRS channel was determined using an anchor-based probabilistic conversion atlas (Tsuzuki et al., 2012). The fNIRS data were then converted to NifTI files using xjView (https://www.alivelearn.net/xjview) and projected onto the cortical surface using BrainNet Viewer (http://www.nitrc.org/projects/bnv/) (Xia et al., 2013).

### Pre-experimental surveys

Prior to the start of the experiment, participants completed the following behavioral questionnaires:

#### Aggression Questionnaire (AQ; Buss and Perry, 1992)

The AQ is a self-assessment questionnaire that measures overall aggression, with subscales for physical aggression, verbal aggression, anger, and hostility. Participants were presented with 29 statements and asked to rate on a 5-point scale the extent to which the statement is characteristic of themselves. Participants’ total score on the AQ was used for analyses. The AQ is widely used as a measure of trait aggression (Archer, 2004; Sekine et al., 2008; da Cunha-Bang et al., 2017).

#### Revised Social Connectedness Scale (SCS-R; Lee *et al*., 2001)

The SCS-R is a 20-item questionnaire that is widely used to measure social connection (Sandstrom and Dunn, 2014; Aknin et al., 2022). Participants were asked to rate the extent to which they agree or disagree with the 20 items on a 6-point scale. Participant’s total score on the SCS-R was used for analyses.

#### Attributional Complexity Scale (ACS; Fletcher *et al*., 1986)

The ACS is a 28-item questionnaire used to measure attributional complexity (i.e. an individual’s tendency and ability to consider complex and multiple causes for their own and others’ behavior). The scale has been externally validated with studies demonstrating that individuals who score highly on attributional complexity spontaneously produced more causes when explaining behavior and selected more complex causal attributions for behavioral events (Fletcher et al., 1986). Attributional complexity has also been shown to be positively correlated with measures of empathy and perspective-taking and negatively correlated with depression and anxiety (Joireman, 2004; Fast et al., 2008). Participant’s total score on the ACS was used for analyses.

### Code and data accessibility

Data and analysis code will be made publicly available upon publication.

## Results

Fifty-eight participants were scanned using fNIRS as they listened to twenty-one hypothetical scenarios in which a character acted in a manner that led to a negative outcome for the participant (total duration: 13 min 30 s; Fig. 1A). After each scenario, participants rated the character’s behavior on hostility, intentionality, and blameworthiness, as well as how angry they would feel if the incident had actually happened. Within a participant, hostility ratings were correlated with intentionality (average *r* = 0.82, SE = 0.018, *t*(58) = 46.47, *p* < 0.001), blameworthiness (average *r* = 0.60, SE = 0.026, *t*(58) = 22.98, *p* < 0.001) and anger ratings across scenarios (average *r* = 0.41, SE = 0.029, *t*(58) = 14.32, *p* < 0.001), suggesting that participants were more likely to attribute intentionality and blame to characters they rated as hostile, and would also be more angry at those characters. Across participants, average hostility ratings were correlated with average intentionality (*r* = 0.91, *t*(58) = 16.7, *p* < 0.001), blameworthiness (*r* =0.76, *t*(58) = 8.99, *p* < 0.001) and anger ratings (*r* = 0.81, *t*(58) = 10.51, *p* < 0.001), indicating that the participants who were more likely to perceive the characters as acting in a hostile manner were the same participants who were likely to attribute blame and intentionality to the characters, as well as experience greater anger in these situations. For each participant, we computed the average hostility ratings across all scenarios as an individual difference measure of hostile attribution bias (Fig. 1B).

### Hostile attribution bias shapes neural synchrony in the ventromedial prefrontal cortex

We employed intersubject representational similarity analysis (IS-RSA; Nguyen *et al*., 2019; Finn *et al*., 2020; Fig. 2A) to identify neural correlates of hostile attribution bias. The intuition behind the analytical approach is that participants with similar levels of hostile attribution bias would process the narratives in a similar fashion, and brain regions associated with hostile attribution bias would exhibit similar temporal dynamics while listening to the narratives. For each fNIRS channel of each participant, we extracted and concatenated the activity time course for the 21 narratives, and z-scored the time courses across time. These time courses capture the fluctuations in participants’ neural activity as they listened to the narratives. For each channel, we then computed a *neural distance matrix* by computing the 1 - Pearson correlation between the time courses of every pair of participants. The neural distance matrix for a given channel reflects the similarity structure in temporal dynamics across participants, with smaller distances denoting pairs of participants with similar neural responses to the stories.

To compute a corresponding *behavioral distance matrix*, we calculated the absolute difference in hostile attribution bias scores between every pair of participants. The behavioral distance matrix reflects the similarity structure in hostile attribution bias scores across participants, with smaller distances denoting pairs of participants with similar levels of hostile attribution bias. For each fNIRS channel, similarity between the neural and behavioral distance matrices was then measured as the Pearson correlation between the lower triangles of the two matrices. The resulting *r*-value would thus reflect the extent to which similarity in hostile attribution bias tracks similarity in neural responses. Statistical significance of the *r*-value was assessed non-parametrically using a Mantel test (Mantel, 1967; Kriegeskorte *et al*., 2008; see Methods), and corrected for multiple comparisons across 20 channels by controlling for the false discovery rate (FDR) of *q* < 0.05.

After controlling for multiple comparisons, this analysis yielded a single significant channel corresponding to the left ventromedial prefrontal cortex (channel 11: VMPFC; *r* = 0.104, *p* = 0.001, *q* = 0.024; Fig. 2B; Table 2-1). The same relationship was not observed in the right VMPFC (channel 13; *r* = 0.005, *p* = 0.415, *q* = 0.792). As a test of whether our results were robust to a different metric of neural distance, we repeated the analysis with the neural distance matrix constructed from the Euclidean distance in channel time course between pairs of participants (Fig. S1A). This yielded identical results, where the channel corresponding to the left VMPFC was the only channel where similarity in levels of hostile attribution bias correlated with similarity in neural responses (*r* = 0.104, *p* < 0.001, *q* = 0.014; Fig. S1B; Table 2-2). The VMPFC has been previously shown to encode subjective evaluations and appraisals (Hutcherson et al., 2015; Chang et al., 2021). Here, our results extend these earlier findings by providing evidence that the subjective interpretation of social situations is shaped by hostile attribution bias and encoded in temporal dynamics of VMPFC activity.

In a series of exploratory analyses, we examined the relationship between neural synchrony and intersubject similarity in intentionality attributions, anger and assessments of blameworthiness. We reran the IS-RSA with behavioral distance matrices constructed from intentionality, anger and blameworthiness ratings. Similar results were observed with intentionality ratings, in that pairwise distances in intentionality ratings between participants were correlated with intersubject dissimilarity in left VMPFC activity time courses (*r* = 0.095, *p* = 0.002, *q* = 0.036; Table 2-3). In contrast, no significant channels were observed with anger and blameworthiness ratings (all *q*s > 0.10; Table 2-3). These results suggest that neural synchrony in the left VMPFC was shaped by shared interpretations of hostile intentions, rather than by shared negative affect due to the negative actions of the characters.

We also considered alternative models of the relationship between neural similarity and levels of hostile attribution bias. We constructed a behavioral distance matrix where participants with low levels of hostile attribution bias were similar to other participants with low levels of hostile attribution bias and less similar to participants with high levels of hostile attribution bias, but participants with high levels of hostile attribution bias were idiosyncratic (i.e. distinct from both other participants with high hostile attribution bias and participants with low levels of hostile attribution bias). This model allows us to test for what is often referred to as the *Anna Karenina* (or *AnnaK*) principle (Finn et al., 2020; Baek et al., 2023). Here, a positive correlation between the AnnaK behavioral matrix and the neural distance matrix would indicate a channel where the neural responses were similar between participants with low levels of hostile attribution bias and idiosyncratic for participants with high levels of hostile attribution bias. In contrast, a negative correlation would indicate a channel where the neural responses were similar between participants with high levels of hostile attribution bias and idiosyncratic for participants with low levels of hostile attribution bias. We found no significant correlations across the 20 channels (all *q*s > 0.30), indicating that the AnnaK models did not provide a good account of the relationship between hostile attribution bias and neural responses.

### Divergence in VMPFC synchrony was driven by narratives with ambiguous intent

The 21 audio narratives can be categorized into scenarios that on average elicit hostile, benign or ambiguous attributions of the character’s intentions (Epps and Kendall, 1995). To investigate if these scenarios are processed differently, we ran the IS-RSA separately for hostile, benign and ambiguous narratives. When the analysis was applied to data from the ambiguous narratives, we replicated the IS-RSA results where the left VMPFC (channel 11) was the only channel for which the neural distance correlated significantly with the absolute difference in hostile attribution bias scores (*r* = 0.1143, *p* < 0.001, *q* = 0.0140).

In contrast, the IS-RSA yielded no significant channels for hostile and benign narratives (all *q*s > 0.10). In other words, neural activity diverged between participants with different levels of hostile attribution bias when listening to narratives where the intent of the social other was ambiguous, but not when listening to narratives where the intent was clearly hostile or benign. One interpretation of these results is that narratives where social intentions were clearly hostile or benign left less room for divergent interpretations, such that participants were likely to make similar attributions regardless of individual differences in hostile attribution bias. However, when intentions were ambiguous, participants’ interpretations were more strongly shaped by their intrinsic predispositions to attribute hostile or benign intentions, thus neural responses diverged between participants with varying levels of hostile attribution bias. This interpretation is consistent with several studies which have suggested that it is in ambiguous situations, where situational cues are lacking, that hostile attribution bias exerts the strongest influence (Combs et al., 2007; Pinkham et al., 2014; Neumann et al., 2020, 2021). Consequently, ambiguous scenarios may provide sufficient information for evaluating hostile attribution bias.

### Synchrony-based classification of hostile attribution bias

Our results indicate that neural activity diverged between individuals with varying levels of hostile attribution bias. Could we then identify participants with high or low levels of hostile attribution bias based on their neural responses to the narratives? We adapted a synchrony-based classification approach used in prior fNIRS and fMRI studies (Fig. 3A; Yeshurun et al., 2017; Dieffenbach et al., 2021) to test this possibility. We first grouped participants into those with high and low levels of hostile attribution bias based on a median split of their hostile attribution scores. Following a leave-one-out cross-validation approach, we held-out the data of one participant. For the remaining participants, we averaged the neural time courses of each channel separately for the participants with high and low hostile attribution bias. These average neural time courses can be thought of as a “template” time course reflecting how the average high and low hostile attribution bias participant processed the 21 narratives. We then computed the correlation between the time course of the held-out participant and each of the two template time courses. If there is a stronger correlation with the high hostile attribution bias template, we classified the participant as having high hostile attribution bias; and if there is a stronger correlation with the low hostile attribution bias template, we classified the participants as having low hostile attribution bias.

**Figure 3.**
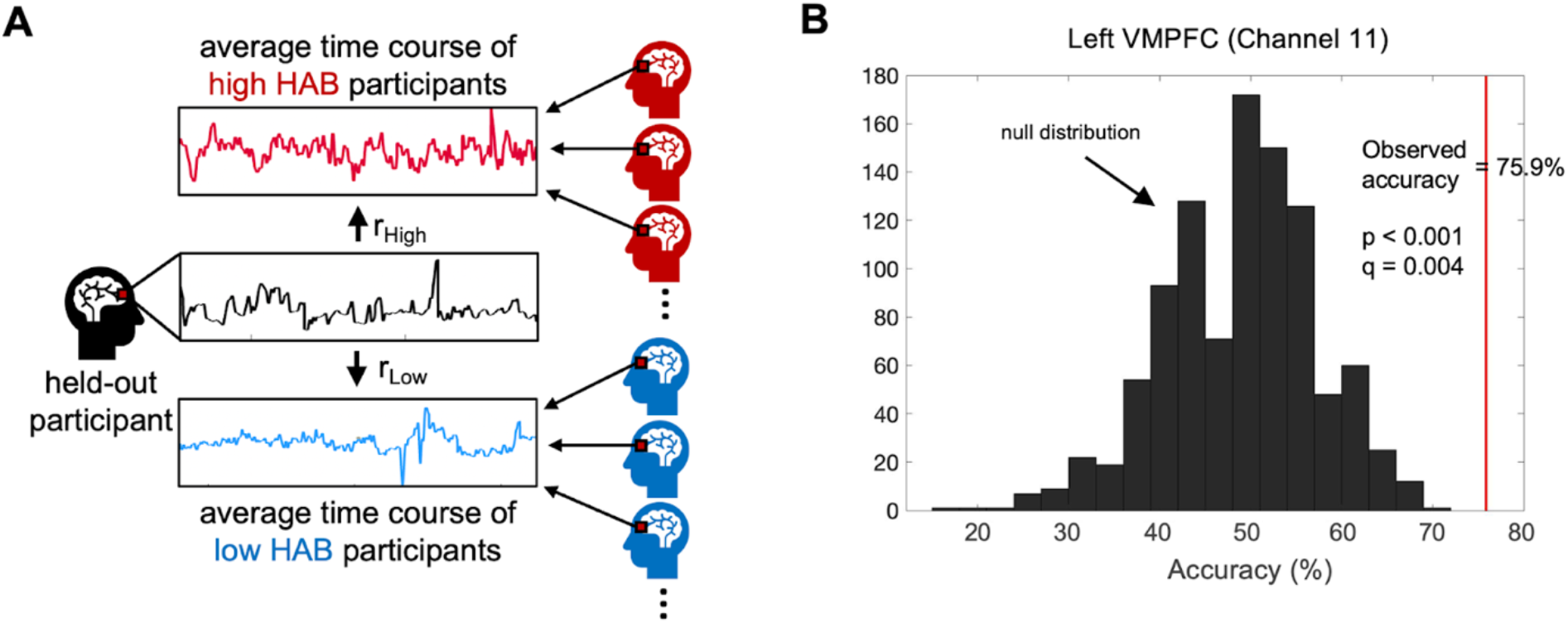
Synchrony-based classification of hostile attribution bias. **(A)** Schematic of classification approach. Following a leave-one-out cross-validation approach, we correlated the activity time course of a held-out participant with the average time course of participants with high and low hostile attribution bias. The held-out participant was classified as the group whose time course they were more correlated to. **(B)** Classification accuracy in the Left VMPFC (Channel 11). Red line indicates observed classification accuracy. Histogram shows null distribution generated from a non-parametric permutation test.

Classification accuracy was significantly above chance in the left VMPFC (channel 11: accuracy = 75.86%, *p* < 0.001, *q* = 0.004; Fig. 3B; Table 3-1), indicating that time courses were sufficiently consistent within a group and distinct between groups such that we were able to distinguish between participants with high and low hostile attribution bias. Classification accuracy was also significantly above chance in the left dorsomedial prefrontal cortex (DMPFC; channel 8: accuracy = 65.5%, *p* = 0.0167, *q* = 0.167), a region that we and others have shown to also be associated with subjective interpretations of narrative information (Yeshurun et al., 2017; Finn et al., 2018; Leong et al., 2020; Dieffenbach et al., 2021), though this result did not survive correction for multiple comparisons.

### No significant differences in mean neural activity between hostile, ambiguous and benign narratives

Thus far, our analyses have focused on the temporal dynamics in neural activity during narrative listening. We next examined whether mean activity differed between hostile, ambiguous and benign narratives, as well as whether differences between narrative types would be moderated by individual differences in hostile attribution bias. For each participant and each channel, we calculated the mean activity over the course of each scenario, and averaged the mean activity separately for hostile, ambiguous and benign narratives. Across participants, mean neural activity was not significantly different between the three types of narratives (all *q*s > 0.05; Fig. 4A depicts results for left VMPFC; see Table 4-1 for results for all channels). Even though there were no significant differences in mean neural activity between narrative types, it was possible that the mean neural responses to the narrative types depended on participants’ level of hostile attribution bias. For example, relative to a participant with low hostile attribution bias, a participant with high hostile attribution bias might have stronger responses to hostile narratives than benign narratives. To test this possibility, we computed the correlation between participants’ hostile attribution bias scores and the difference in mean neural activity between narrative types. Across the three contrasts, difference in mean neural activity was not correlated with hostile attribution bias scores (all *q*s > 0.10; Table 4-2). Taken together, these results suggest that differences in how participants processed hostile, ambiguous and benign narratives were unlikely to be reflected in the mean neural activity during narrative listening.

**Figure 4.**
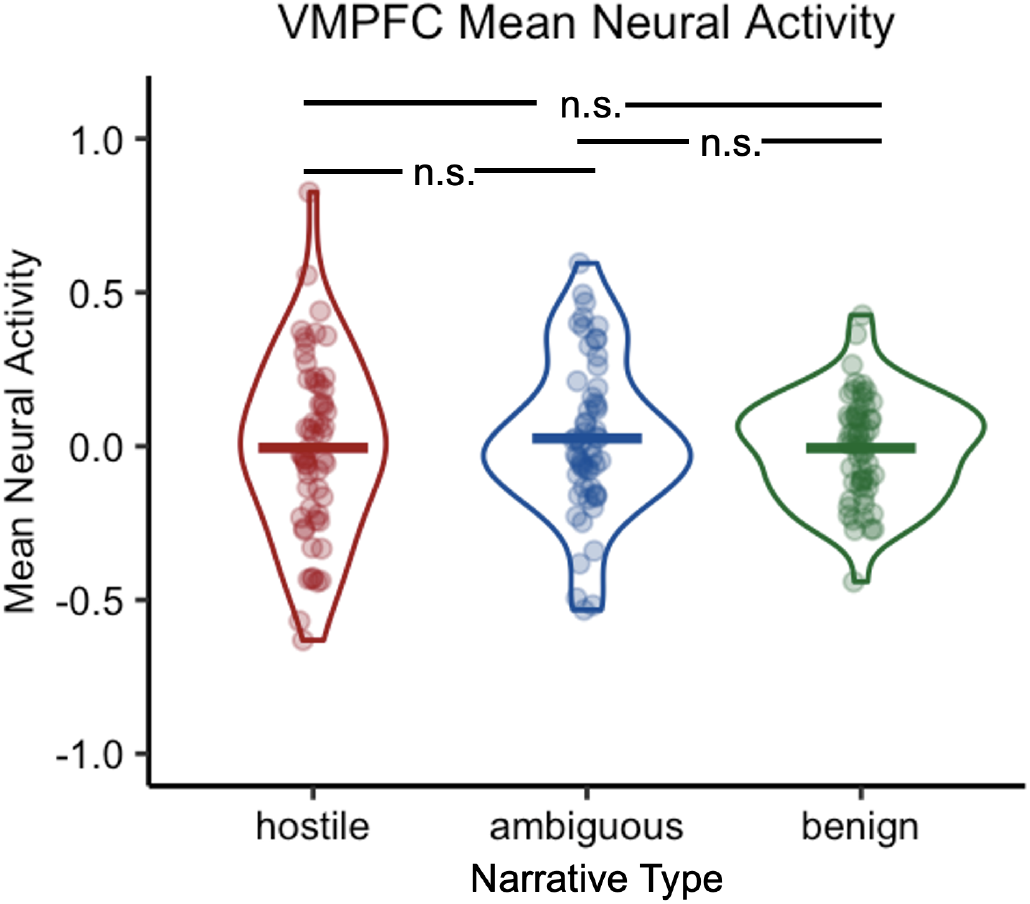
Violin plots showing participants’ mean neural activity in VMPFC during hostile, ambiguous and benign stories. No significant difference was observed between the narrative types. Each data point indicates an individual participant. Thick solid line indicates the corresponding group mean.

### Attributional complexity is negatively correlated with hostile attribution bias

Understanding the predictors of hostile attribution bias can reveal the underlying psychological factors and might allow researchers to develop interventions that mitigate the bias. Here, we consider two potential predictors: attributional complexity and social connectedness. Attributional complexity was measured using the Attributional Complexity Scale (Fletcher et al., 1986), and refers to one’s tendency and ability to consider complex and multiple causes for their own and others’ behavior, taking into account both external (i.e. situational) and internal (i.e. dispositional) factors. On one hand, attributional complexity might exacerbate hostile attribution bias as the individual might be more likely to generate complex explanations to rationalize their hostile attributions. On the other hand, attributional complexity might buffer against hostile attribution bias by encouraging a more nuanced understanding of others’ behavior. Here, we found that attributional complexity was negatively correlated with hostile attribution bias (*r* = −0.314, *t*(58) = −2.52, *p* = 0.0146; Fig. 5), suggesting that individuals who tend to infer complex attributions were less likely to make hostile attributions.

**Figure 5.**
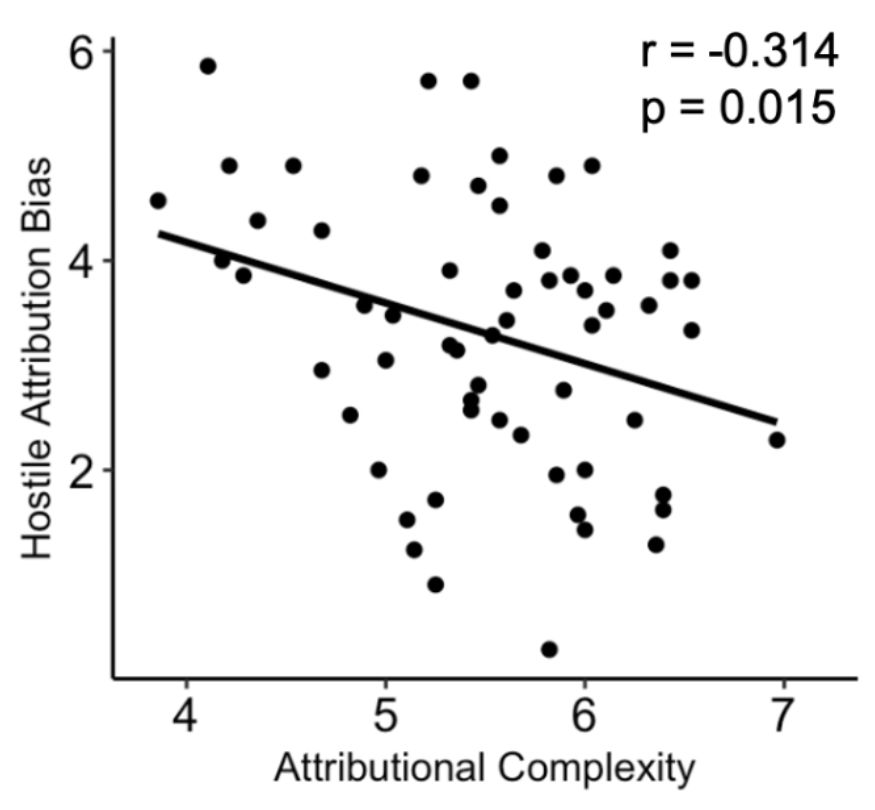
Attributional complexity is negatively correlated with hostile attribution bias. Attributional complexity refers to one’s tendency and ability to consider complex and multiple causes for their own and others’ behavior. Each data point indicates an individual participant.

Social connectedness, as measured using the revised Social Connectedness Scale (SCS-R; Lee *et al*., 2001) was not correlated with hostile attribution bias (*r* = −0.12, *t*(58) = −0.95, *p* = 0.346). As aggression is often cited as a downstream consequence of hostile attributions (Neumann et al., 2015; Coccaro et al., 2016; Klein Tuente et al., 2019), we sought to test the relationship between aggression and hostile attribution bias. In our sample, trait aggression, as measured using the Aggression Questionnaire (AQ; Buss and Perry, 1992), was only marginally correlated with hostile attribution bias (*r* = 0.238, *t*(58) = 1.87, *p* = 0.067), possibly because our sample consisted of healthy participants in a college community while many of the prior studies have recruited special populations (e.g., patients with clinical diagnosis, incarcerated individuals).

## Discussion

In this study, we combined fNIRS and a narrative listening paradigm to examine how neural activity during the processing of social situations is influenced by individual differences in hostile attribution bias. During narrative listening, temporal dynamics of left VMPFC activity were more synchronous between individuals with similar levels of hostile attribution bias. The effect of hostile attribution bias on neural synchrony was particularly strong during narratives where the intention of the social other was ambiguous, suggesting that uncertain social contexts were more susceptible to the influence of intrinsic attributional biases. Left VMPFC time courses between participants with high and low hostile attribution bias were systematically different such that a synchrony-based classifier could distinguish between the two groups from neural activity alone. We also found that people who infer more complex attributions of others’ behaviors were less likely to make hostile attributions. Together, our results suggest that hostile attribution bias influences subjective interpretations of social situations via differential responses in the left VMPFC.

The VMPFC has been implicated in a broad range of complex cognitive functions, including evaluating the subjective value of economic and moral decisions (Bartra et al., 2013; Hutcherson et al., 2015), inferring traits and intentions from behavior (Young et al., 2010; Leopold et al., 2012; Kestemont et al., 2016), and narrative comprehension (van Kesteren et al., 2010; Burin et al., 2014; Yeshurun et al., 2017). One proposed account that unifies these disparate findings is that the VMPFC integrates information about the external world with internal states and prior beliefs to generate “affective meaning” - the subjective appraisal of a situation, experience or object, including its relevance to oneself and the affective states it engenders (Roy et al., 2012; Chang et al., 2021). In line with this proposal, a recent fMRI study by Chang and colleagues (2021) mapped patterns of activity in the VMPFC onto specific affective states while participants watched a 45-minute episode from a television drama. The temporal dynamics of these patterns were largely idiosyncratic across individuals, consistent with the hypothesis that the VMPFC encodes subjective affective meaning that varies from person-to-person. Our results suggest that temporal dynamics in the VMPFC track with the subjective interpretation of ambiguous social situations, and are in line with these earlier findings. Furthermore, our work extends past work by demonstrating that VMPFC dynamics are not necessarily idiosyncratic, but can be synchronized between individuals with similar socio-cognitive biases. Specifically, our results suggest that hostile attribution bias acts as an intrinsic “prior” that shapes subjective interpretations and VMPFC time courses when processing social information.

We found that hostile attribution bias was associated with neural synchrony in the left, but not the right, VMPFC. Previous studies have documented a similar left-right asymmetry in the prefrontal cortex, with regions in the left hemisphere showing stronger associations with anger, aggression and hostility than corresponding regions in the right hemisphere (Gansler et al., 2009; Harmon-Jones et al., 2010; Wang et al., 2018). For example, larger gray matter volume in the left OFC, a region immediately adjacent to the left VMPFC, was associated with higher levels of hostile attribution bias (Quan et al., 2019). Left frontal activity, as measured using EEG, was higher in participants who had just been insulted or socially rejected, with the magnitude of left frontal response correlating with subsequent aggression (Harmon-Jones and Sigelman, 2001; Verona et al., 2009). Furthermore, stimulating the left frontal cortex using non-invasive brain stimulation increased aggression in a laboratory task, while stimulating the right frontal cortex had no effect (Hortensius et al., 2012). It is worthwhile to note, however, that intersubject similarity in anger and blameworthiness ratings did not predict left VMPFC synchrony in our study. Thus, left VMPFC synchrony did not reflect shared anger at the character in the scenario. Instead, we believe left VMPFC activity was specifically encoding shared interpretations of the character’s hostile intentions. Consistent with this hypothesis, similarity in intentionality ratings was also associated with left VMPFC synchrony.

A meta-analysis of 43 structural and functional neuroimaging studies found that reduced structure and function in the lateral prefrontal cortices (LPFC) was associated with antisocial behavior (e.g., aggression, violence) (Yang and Raine, 2009). Furthermore, increased LPFC activity following negative social interactions is associated with decreases in retaliatory aggression (Achterberg et al., 2016, 2020), and disrupting LPFC function using non-invasive brain stimulation prior to negative interactions increases retaliatory aggression (Perach-Barzilay et al., 2013; Choy et al., 2018). In our study, we did not find that the LPFC was differentially engaged depending on participants’ level of hostile attribution bias. One possible explanation is that the LPFC is involved in inhibiting aggressive tendencies following negative social interactions, consistent with its role in behavioral inhibition more broadly (Shackman et al., 2009; Aron et al., 2014). In the earlier studies, participants had the opportunity to retaliate against the individual who caused them harm, and the LPFC is engaged to inhibit aggressive behavior. Participants in our task, however, were not able to respond to the hypothetical characters in the scenarios. Instead, the task primed them to consider the intentions of the characters. Without a behavioral response to inhibit, the LPFC is less likely to be engaged, and consequently, the time course of LPFC activity was not modulated by individual differences in hostile attribution bias.

Given the extensive research implicating the prefrontal cortex in subjective appraisals, social cognition and aggression, our study focused specifically on the prefrontal cortex to better understand its role in the manifestation of hostile attribution bias. Our results highlight the role of the VMPFC in encoding subjective interpretations of social situations that were biased by individual differences in hostile attribution bias. We note that there are likely other brain regions involved in this process, including the amygdala, a region often associated with threat detection and processing (Öhman, 2005; Coccaro et al., 2007), which also has dense reciprocal connections with the VMPFC (Barbas, 2000; Kim et al., 2011). Unfortunately, fNIRS is limited to measuring cortical activity near the surface of the brain and is unable to reach deep brain structures such as the amygdala. A whole-brain functional neuroimaging method (e.g., fMRI) will be needed to examine if the amygdala-VMPFC interactions contribute to hostile attribution bias. Another candidate region of interest is the temporoparietal junction (TPJ), which is widely thought to be involved in reasoning about others’ intentions and beliefs (Saxe and Kanwisher, 2003; Van Overwalle, 2009). Unlike the amygdala, the TPJ is located closer to the cortical surface (Jiang et al., 2015), and future studies can use fNIRS to examine the relationship between TPJ activity and hostile attribution bias.

In our study, participants who scored higher on attributional complexity exhibited less hostile attribution bias in their behavioral responses to the narratives, a relationship which to our knowledge has not been previously shown. This result suggests that individuals with a propensity to consider multifaceted explanations for behavior are less inclined to interpret ambiguous behaviors as malicious in intent. If so, fostering attributional complexity could be a potential strategy to mitigate hostile attribution bias and ultimately promote healthier social interactions. In line with this hypothesis, a recent study with individuals with traumatic brain injury found that a group-based intervention that promotes thinking about alternative causes of behavior and perspective-taking in simulated and experienced social situations was effective at reducing attributions of intent, blame, anger and aggression responses to the same 21 scenarios used in the current study (Neumann et al., 2023). Future work could combine the intervention with neural measures to determine its neural basis, which would inform efforts at developing neuroscience-informed markers of hostile attribution bias and intervention efficacy.

Social interaction involves the decoding and interpretation of subtle cues, the recognition of intent, and the formation of appropriate responses based on those interpretations. Individuals with hostile attribution bias have a distorted perception of social intentions, often resulting in unnecessary misunderstandings and conflicts. Here, our results highlight the influence of hostile attribution bias on shaping the neural responses to social situations, and demonstrate the viability of using fNIRS to investigate the interplay between neural processes and cognitive biases in social perception. As a method, fNIRS offers a unique combination of non-invasiveness, ease of use, and sufficient spatial resolution to localize brain responses to specific regions of the cortex (Yücel et al., 2017). The relative cost-effectiveness of fNIRS makes it an ideal tool for large-scale studies that can more precisely examine individual differences in socio-cognitive biases, as well as the development of scalable tools for use in clinical settings. Additionally, the portability of fNIRS allows for more naturalistic experimental designs that could capture how individual biases interact with social perception in more ecologically valid settings (e.g., an unconstrained conversation). Our study thus paves the way for future research employing fNIRS to better understand the dynamics of social cognition, including how they can break down in the presence of maladaptive biases. This approach holds significant potential to enhance our understanding of the neural processes underlying social cognition and to mitigate the adverse effects of socio-cognitive biases on interpersonal relationships.

## Acknowledgements

We would like to thank the members of the Motivation and Cognition Neuroscience Laboratory and the University of Chicago fNIRS user group for helpful comments on the study, as wekk as the University of Chicago Neuroscience Institute Shared Equipment Award for providing the fNIRS device.

