## supplemental material for "Hostile attribution bias shapes neural synchrony in the left ventromedial prefrontal cortex during ambiguous social narratives"

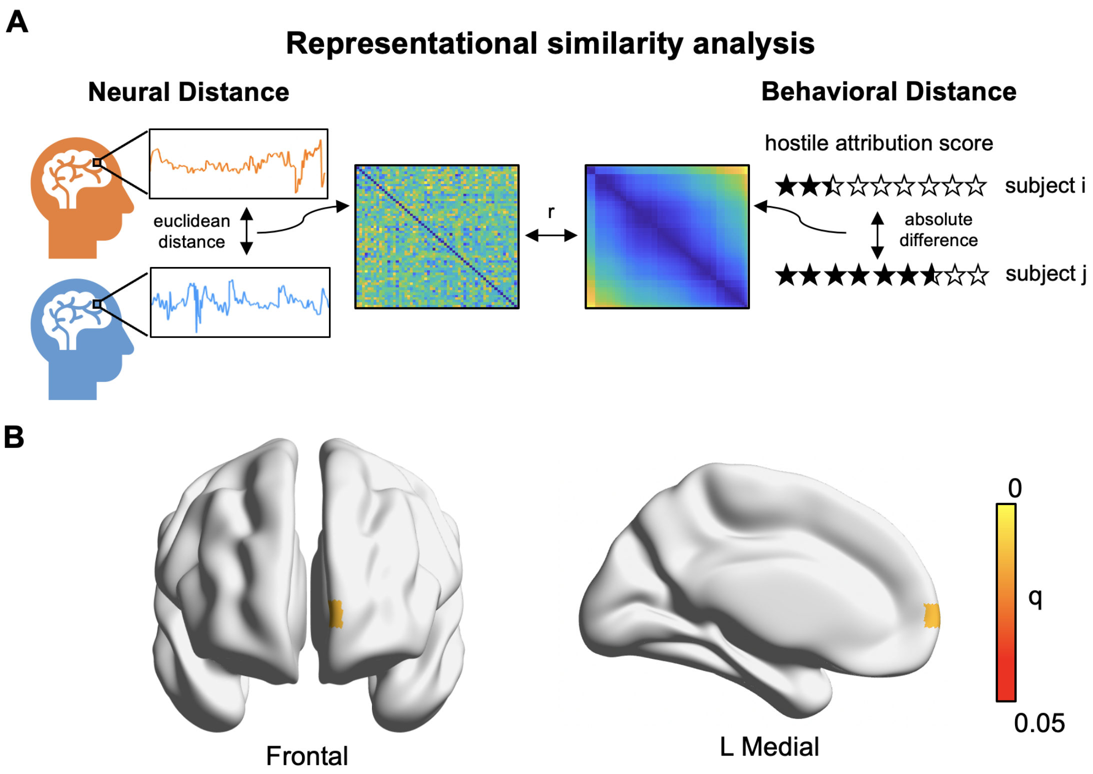


**Figure 2-1. Neural synchrony results replicate using Euclidean distance as measure of neural distance. (A)** Schematic of intersubject-representational similarity analysis. For each channel, we constructed a neural distance matrix by calculating the Euclidean distance between the time courses of every pair of participants. Each neural distance matrix was correlated with a behavioral distance matrix constructed from the absolute difference in hostile attribution bias scores for every pair of participants. **(B)** Left VMPFC (Channel 11) activity time course was more correlated between individuals with similar levels of hostile attribution bias. P values were computed nonparametrically using a Mantel test, and thresholded at a FDR of $q < 0.05$.

**Table 2-1. Intersubject representational similarity analysis results when computing neural distance matrices using correlation distance.** * q < 0.05

| **Channel** | ***r*** | ***p-value*** | ***FDR q-value*** |
|---|---|---|---|
| 1 | -0.030 | 0.8061 | 0.893 |
| 2 | -0.000 | 0.4739 | 0.792 |
| 3 | 0.039 | 0.1417 | 0.733 |
| 4 | 0.011 | 0.3225 | 0.792 |
| 5 | -0.006 | 0.5790 | 0.792 |
| 6 | -0.010 | 0.6241 | 0.792 |
| 7 | -0.062 | 0.9976 | 0.998 |
| 8 | 0.029 | 0.1589 | 0.733 |
| 9 | -0.003 | 0.5235 | 0.792 |
| 10 | -0.006 | 0.5662 | 0.792 |
| 11 | 0.104 | 0.0010 | 0.024* |
| 12 | 0.013 | 0.3372 | 0.792 |
| 13 | 0.005 | 0.4153 | 0.792 |
| 14 | 0.003 | 0.1516 | 0.733 |
| 15 | 0.026 | 0.1863 | 0.733 |
| 16 | 0.022 | 0.2199 | 0.733 |
| 17 | -0.024 | 0.7427 | 0.874 |
| 18 | -0.042 | 0.8487 | 0.893 |
| 19 | -0.005 | 0.5536 | 0.792 |
| 20 | -0.015 | 0.6366 | 0.792 |

**Table 2-2. Intersubject representational similarity analysis results when computing neural distance matrices using Euclidean distance.** * q < 0.05

| **Channel** | ***r*** | ***p-value*** | ***FDR q-value*** |
|---|---|---|---|
| 1 | -0.033 | 0.8255 | 0.869 |
| 2 | 0.001 | 0.4716 | 0.806 |
| 3 | 0.040 | 0.1403 | 0.722 |
| 4 | 0.012 | 0.3109 | 0.806 |
| 5 | -0.005 | 0.5550 | 0.806 |
| 6 | -0.012 | 0.6472 | 0.809 |
| 7 | -0.062 | 0.9981 | 0.998 |
| 8 | 0.030 | 0.1443 | 0.722 |
| 9 | -0.002 | 0.5024 | 0.806 |
| 10 | -0.004 | 0.5384 | 0.806 |
| 11 | 0.104 | 0.0007 | 0.014* |
| 12 | 0.014 | 0.3478 | 0.806 |
| 13 | 0.005 | 0.4125 | 0.806 |
| 14 | 0.032 | 0.1335 | 0.722 |
| 15 | 0.025 | 0.1947 | 0.779 |
| 16 | 0.019 | 0.2461 | 0.806 |
| 17 | -0.024 | 0.7380 | 0.869 |
| 18 | -0.038 | 0.8167 | 0.869 |
| 19 | -0.005 | 0.5639 | 0.806 |
| 20 | -0.015 | 0.6373 | 0.809 |

**Table 2-3. Intersubject representational similarity analysis results correlating neural distance matrix with dissimilarity in intentionality, anger and blameworthiness ratings.** FDR-correction was applied separately for each behavioral measure. * q < 0.05

| Channel | Intentionality | Anger | Blameworthiness |
|---|---|---|---|
| **1** | *r* = -0.016, *p* = 0.672, q = 0.827 | *r* = -0.050, *p* = 0.930, q = 0.930 | *r* = -0.031, *p* = 0.793, q = 0.830 |
| **2** | *r* = 0.009, *p* = 0.356, q = 0.726 | *r* = -0.031, *p* = 0.887, q = 0.930 | *r* = -0.017, *p* = 0.729, q = 0.810 |
| **3** | *r* = 0.025, *p* = 0.247, q = 0.726 | *r* = 0.029, *p* = 0.217, q = 0.609 | *r* = 0.045, *p* = 0.125, q = 0.472 |
| **4** | *r* = -0.003, *p* = 0.534, q = 0.827 | *r* = -0.009, *p* = 0.617, q = 0.822 | *r* = 0.011, *p* = 0.325, q = 0.541 |
| **5** | *r* = 0.014, *p* = 0.296, q = 0.726 | *r* = 0.041, *p* = 0.073, q = 0.609 | *r* = -0.027, *p* = 0.830, q = 0.830 |
| **6** | *r* = 0.033, *p* = 0.125, q = 0.623 | *r* = 0.025, *p* = 0.182, q = 0.609 | *r* = 0.073, *p* = 0.009, q = 0.180 |
| **7** | *r* = -0.039, *p* = 0.950, q = 0.950 | *r* = -0.021, *p* = 0.806, q = 0.930 | *r* = -0.015, *p* = 0.721, q = 0.810 |
| **8** | *r* = 0.046, *p* = 0.059, q = 0.590 | *r* = 0.023, *p* = 0.212, q = 0.609 | *r* = 0.026, *p* = 0.187, q = 0.472 |
| **9** | *r* = -0.018, *p* = 0.728, q = 0.827 | *r* = 0.008, *p* = 0.366, q = 0.609 | *r* = -0.007, *p* = 0.583, q = 0.777 |
| **10** | *r* = 0.007, *p* = 0.385, q = 0.726 | *r* = 0.011, *p* = 0.338, q = 0.609 | *r* = -0.012, *p* = 0.643, q = 0.804 |
| **11** | *r* = 0.095, *p* = 0.002, q = 0.036* | *r* = -0.004, *p* = 0.533, q = 0.761 | *r* = 0.010, *p* = 0.352, q = 0.541 |
| **12** | *r* = 0.008, *p* = 0.399, q = 0.726 | *r* = 0.013, *p* = 0.344, q = 0.609 | *r* = 0.047, *p* = 0.120, q = 0.472 |
| **13** | *r* = -0.019, *p* = 0.744, q = 0.827 | *r* = -0.037, *p* = 0.924, q = 0.930 | *r* = -0.007, *p* = 0.572, q = 0.777 |
| **14** | *r* = -0.004, *p* = 0.540, q = 0.827 | *r* = 0.002, *p* = 0.457, q = 0.702 | *r* =0.016, *p* = 0.277, q = 0.504 |
| **15** | *r* = 0.036, *p* = 0.118, q = 0.623 | *r* = 0.043, *p* = 0.080, q = 0.609 | *r* = 0.026, *p* = 0.189, q = 0.472 |
| **16** | *r* = 0.010, *p* = 0.342, q = 0.726 | *r* = 0.016, *p* = 0.277, q = 0.609 | *r* = 0.038, *p* = 0.110, q = 0.472 |
| **17** | *r* = 0.011, *p* = 0.372, q = 0.726 | *r* = 0.032, *p* = 0.188, q = 0.609 | *r* = 0.037, *p* = 0.158, q = 0.472 |
| **18** | *r* = -0.010, *p* = 0.583, q = 0.827 | *r* = 0.034, *p* = 0.211, q = 0.609 | *r* = 0.030, *p* = 0.248, q = 0.495 |
| **19** | *r* = -0.015, *p* = 0.696, q = 0.827 | *r* = 0.016, *p* = 0.274, q = 0.609 | *r* = 0.041, *p* = 0.081, q = 0.472 |
| **20** | *r* = -0.055, *p* = 0.906, q = 0.950 | *r* = -0.027, *p* = 0.730, q = 0.912 | *r* = 0.034, *p* = 0.218, q = 0.485 |

**Table 3-1. Single-channel classification accuracy.** * q < 0.01

| **Channel** | ***Accuracy*** | ***p-value*** | ***FDR q-value*** |
|---|---|---|---|
| 1 | 0.569 | 0.110 | 0.442 |
| 2 | 0.483 | 0.507 | 0.643 |
| 3 | 0.517 | 0.339 | 0.643 |
| 4 | 0.500 | 0.515 | 0.643 |
| 5 | 0.586 | 0.078 | 0.401 |
| 6 | 0.535 | 0.339 | 0.643 |
| 7 | 0.535 | 0.350 | 0.643 |
| 8 | 0.655 | 0.017 | 0.167 |
| 9 | 0.466 | 0.047 | 0.643 |
| 10 | 0.500 | 0.425 | 0.643 |
| 11 | 0.759 | < 0.001 | 0.004** |
| 12 | 0.621 | 0.080 | 0.401 |
| 13 | 0.535 | 0.385 | 0.643 |
| 14 | 0.517 | 0.387 | 0.643 |
| 15 | 0.483 | 0.552 | 0.649 |
| 16 | 0.483 | 0.618 | 0.687 |
| 17 | 0.517 | 0.339 | 0.643 |
| 18 | 0.397 | 0.820 | 0.820 |
| 19 | 0.535 | 0.380 | 0.643 |
| 20 | 0.448 | 0.722 | 0.760 |

**Table 4-1. Differences in mean activity between hostile, ambiguous and benign narratives.** M indicates mean difference between two narrative types. FDR-correction was applied separately for each contrast.

| Channel | hostile - ambiguous | hostile - benign | ambiguous - benign |
|---|---|---|---|
| **1** | M = 0.07, $t(53) = 2.23$, $p = 0.030$, $q = 0.244$ | M = -0.07, $t(53) = -1.65$, $p = 0.106$, $q = 0.545$ | M = -0.14, $t(53) = -3.02$, $p = 0.004$, $q = 0.078$ |
| **2** | M = 0.08, $t(54) = 2.14$, $p = 0.037$, $q = 0.244$ | M = 0.05, $t(54) = 1.00$, $p = 0.322$, $q = 0.545$ | M = -0.03, $t(54) = -0.57$, $p = 0.569$, $q = 0.711$ |
| **3** | M = 0.01, $t(54) = 0.22$, $p = 0.829$, $q = 0.921$ | M = 0.06, $t(54) = 1.61$, $p = 0.114$, $q = 0.545$ | M = 0.06, $t(54) = 1.35$, $p = 0.183$, $q = 0.523$ |
| **4** | M = 0.04, $t(56) = 1.27$, $p = 0.210$, $q = 0.700$ | M = 0.07, $t(56) = 1.66$, $p = 0.103$, $q = 0.545$ | M = 0.03, $t(56) = 0.77$, $p = 0.442$, $q = 0.632$ |
| **5** | M = 0.09, $t(53) = 2.31$, $p = 0.025$, $q = 0.244$ | M = 0.04, $t(53) = 0.99$, $p = 0.325$, $q = 0.545$ | M = -0.05, $t(53) = -1.18$, $p = 0.243$, $q = 0.540$ |
| **6** | M = 0.05, $t(56) = 1.36$, $p = 0.179$, $q = 0.700$ | M = 0.00, $t(56) = -0.17$, $p = 0.868$, $q = 0.868$ | M = -0.06, $t(56) = -1.45$, $p = 0.153$, $q = 0.510$ |
| **7** | M = 0.01, $t(56) = 0.28$, $p = 0.782$, $q = 0.920$ | M = -0.07, $t(56) = -1.63$, $p = 0.108$, $q = 0.545$ | M = -0.07, $t(56) = -1.59$, $p = 0.118$, $q = 0.471$ |
| **8** | M = 0.07, $t(55) = 1.92$, $p = 0.060$, $q = 0.301$ | M = 0.03, $t(55) = 0.95$, $p = 0.346$, $q = 0.545$ | M = -0.03, $t(55) = -0.85$, $p = 0.398$, $q = 0.632$ |
| **9** | M = 0.02, $t(51) = 0.39$, $p = 0.696$, $q = 0.870$ | M = -0.02, $t(51) = -0.59$, $p = 0.560$, $q = 0.701$ | M = -0.04, $t(51) = -0.92$, $p = 0.364$, $q = 0.632$ |
| **10** | M = 0.03, $t(54) = 0.94$, $p = 0.351$, $q = 0.829$ | M = -0.04, $t(54) = -1.32$, $p = 0.192$, $q = 0.545$ | M = -0.08, $t(54) = -1.86$, $p = 0.068$, $q = 0.471$ |
| **11** | M = 0.03, $t(56) = 0.90$, $p = 0.373$, $q = 0.829$ | M = 0.03, $t(56) = 0.77$, $p = 0.444$, $q = 0.592$ | M = -0.01, $t(56) = -0.14$, $p = 0.889$, $q = 0.889$ |
| **12** | M = 0.02, $t(57) = 0.40$, $p = 0.688$, $q = 0.870$ | M = -0.01, $t(57) = -0.34$, $p = 0.733$, $q = 0.814$ | M = -0.03, $t(57) = -0.60$, $p = 0.548$, $q = 0.711$ |
| **13** | M = 0.04, $t(57) = 1.02$, $p = 0.312$, $q = 0.829$ | M = 0.05, $t(57) = 1.15$, $p = 0.257$, $q = 0.545$ | M = 0.01, $t(57) = 0.32$, $p = 0.750$, $q = 0.868$ |
| **14** | M = 0.02, $t(55) = 0.66$, $p = 0.510$, $q = 0.870$ | M = -0.04, $t(55) = -0.94$, $p = 0.354$, $q = 0.545$ | M = -0.07, $t(55) = -1.66$, $p = 0.102$, $q = 0.471$ |
| **15** | M = 0.00, $t(55) = 0.03$, $p = 0.975$, $q = 0.975$ | M = -0.03, $t(55) = -0.94$, $p = 0.351$, $q = 0.545$ | M = -0.04, $t(55) = -0.81$, $p = 0.421$, $q = 0.632$ |
| **16** | M = 0.00, $t(57) = 0.08$, $p = 0.936$, $q = 0.975$ | M = 0.01, $t(57) = 0.37$, $p = 0.714$, $q = 0.814$ | M = 0.01, $t(57) = 0.28$, $p = 0.781$, $q = 0.868$ |
| **17** | M = 0.02, $t(54) = 0.58$, $p = 0.562$, $q = 0.870$ | M = -0.03, $t(54) = -0.87$, $p = 0.388$, $q = 0.554$ | M = -0.05, $t(54) = -1.18$, $p = 0.243$, $q = 0.540$ |
| **18** | M = 0.02, $t(53) = 0.48$, $p = 0.631$, $q = 0.870$ | M = -0.05, $t(53) = -1.22$, $p = 0.228$, $q = 0.545$ | M = -0.07, $t(53) = -1.69$, $p = 0.097$, $q = 0.471$ |
| **19** | M = 0.02, $t(57) = 0.62$, $p = 0.541$, $q = 0.870$ | M = 0.06, $t(57) = 1.35$, $p = 0.181$, $q = 0.545$ | M = 0.04, $t(57) = 0.87$, $p = 0.390$, $q = 0.632$ |
| **20** | M = 0.02, $t(56) =0.44$, $p = 0.663$, $q = 0.870$ | M = 0.01, $t(56) = 0.25$, $p = 0.803$, $q = 0.845$ | M = -0.01, $t(56) = -0.15$, $p = 0.886$, $q = 0.889$ |

**Table 4-2. Correlation between hostile attribution bias scores and difference in mean activity between hostile, ambiguous and benign narratives.** FDR-correction was applied separately for each contrast.

| Channel | hostile - ambiguous | hostile - benign | ambiguous - benign |
|---|---|---|---|
| **1** | *r* = 0.246, *p* = 0.072, q = 1.000 | *r* = 0.261, *p* = 0.056, q = 0.902 | *r* = 0.046, *p* = 0.741, q = 1.000 |
| **2** | *r* = 0.054, *p* = 0.696, q = 1.000 | *r* = 0.279, *p* = 0.039, q = 0.681 | *r* = 0.222, *p* = 0.103, q = 1.000 |
| **3** | *r* = 0.367, *p* = 0.006, q = 0.118 | *r* = 0.300, *p* = 0.026, q = 0.522 | *r* = -0.069, *p* = 0.618, q = 1.000 |
| **4** | *r* = 0.066, *p* = 0.627, q = 1.000 | *r* = 0.175, *p* = 0.194, q = 1.000 | *r* = 0.132, *p* = 0.326, q = 1.000 |
| **5** | *r* = -0.029, *p* = 0.833, q = 1.000 | *r* = -0.007, *p* = 0.960, q = 1.000 | *r* = 0.020, *p* = 0.884, q = 1.000 |
| **6** | *r* = 0.117, *p* = 0.385, q = 1.000 | *r* = 0.155, *p* = 0.251, q = 1.000 | *r* = 0.027, *p* = 0.842, q = 1.000 |
| **7** | *r* = -0.105, *p* = 0.435, q = 1.000 | *r* = -0.085, *p* = 0.528, q = 1.000 | *r* = 0.005, *p* = 0.973, q = 1.000 |
| **8** | *r* = -0.088, *p* = 0.520, q = 1.000 | *r* = 0.068, *p* = 0.621, q = 1.000 | *r* = 0.130, *p* = 0.340, q = 1.000 |
| **9** | *r* = -0.004, *p* = 0.977, q = 1.000 | *r* = 0.076, *p* = 0.590, q = 1.000 | *r* = 0.076, *p* = 0.593, q = 1.000 |
| **10** | *r* = 0.091, *p* = 0.508, q = 1.000 | *r* = 0.281, *p* = 0.038, q = 0.681 | *r* = 0.147, *p* = 0.285, q = 1.000 |
| **11** | *r* = 0.068, *p* = 0.613, q = 1.000 | *r* = 0.284, *p* = 0.032, q = 0.608 | *r* = 0.184, *p* = 0.170, q = 1.000 |
| **12** | *r* = 0.065, *p* = 0.630, q = 1.000 | *r* = 0.197, *p* = 0.139, q = 1.000 | *r* = 0.114, *p* = 0.396, q = 1.000 |
| **13** | *r* = 0.095, *p* = 0.480, q = 1.000 | *r* = 0.241, *p* = 0.068, q = 1.000 | *r* = 0.163, *p* = 0.222, q = 1.000 |
| **14** | *r* = -0.059, *p* = 0.664, q = 1.000 | *r* = 0.045, *p* = 0.745, q = 1.000 | *r* = 0.104, *p* = 0.446, q = 1.000 |
| **15** | *r* = -0.207, *p* = 0.126, q = 1.000 | *r* = -0.227, *p* = 0.093, q = 1.000 | *r* = 0.006, *p* = 0.967, q = 1.000 |
| **16** | *r* = -0.003, *p* = 0.984, q = 1.000 | *r* = 0.091, *p* = 0.497, q = 1.000 | *r* = 0.085, *p* = 0.524, q = 1.000 |
| **17** | *r* = -0.096, *p* = 0.485, q = 1.000 | *r* = -0.189, *p* = 0.167, q = 1.000 | *r* = -0.060, *p* = 0.664, q = 1.000 |
| **18** | *r* = 0.176, *p* = 0.203, q = 1.000 | *r* = 0.032, *p* = 0.816, q = 1.000 | *r* = -0.127, *p* = 0.362, q = 1.000 |
| **19** | *r* = 0.107, *p* = 0.425, q = 1.000 | *r* = 0.183, *p* = 0.170, q = 1.000 | *r* = 0.099, *p* = 0.458, q = 1.000 |
| **20** | *r* = 0.192, *p* = 0.152, q = 1.000 | *r* = 0.195, *p* = 0.146, q = 1.000 | *r* = 0.022, *p* = 0.869, q = 1.000 |
